# *Cutibacterium acnes*-derived short-chain fatty acids drive lipogenesis and induce holocrine secretion in human sebocytes

**DOI:** 10.64898/2026.09.23.753677

**Authors:** Ann Don Low, Omkar S. Kulkarni, Francoise Brillet, Eganathan Kaliyamoorthy, Angeline S.L. Tay, Scott Rice, Viduthalai R. Regina, Tarun Chopra

## Abstract

Sebum lipids are one of the key ecological determinants of skin microbiome composition, where sebaceous sites are enriched with sebum utilising microbes such as *Cutibacterium acnes*. While sebaceous gland (SG) activation and sebogenesis are classically viewed as host-regulated processes, the association of *C. acnes* expansion with sebum production in the skin suggests a possible bidirectional host-microbe regulation. Using bacterial supernatants, we combined image-based lipid quantification, GC-FID metabolite profiling, lipid secretion quantification, and transcriptomic analysis to determine the metabolic drivers of the interaction between *C. acnes* and sebocytes. We found that *C. acnes* secretome significantly increased lipid droplet accumulation in sebocytes and propionate as the primary driver of this lipogenic response. Mechanistically, propionate reprogrammed the central carbon metabolism, redirecting carbon flux toward energy production and generation of precursors for lipid biosynthesis. Propionate also promoted lipid assembly pathways and modulated the composition of the secreted lipids from the treated sebocytes. We further show that propionate promotes expression of late sebocyte differentiation markers associated with holocrine secretion. Together, these findings redefine the relationship between microbiome and sebaceous glands, and identify *C. canes* as an active regulator promoting both sebogenesis and holocrine secretion for the release of lipids in the pilosebaceous unit.

## INTRODUCTION

Human sebaceous skin represents a distinctive microbial habitat, characterized by high lipid availability, and a distinct microbiome, presumable shaped by the high amount of sebum secreted by the sebaceous gland (Picardo et al. 2009; SanMiguel and Grice 2015; Stücker et al. 2002). In particular, *Cutibacterium acnes* (formerly *Propionibacterium acnes*) is a dominant inhabitant of the sebaceous skin colonizing various anatomical structures including hair follicles and pores of the skin (Flowers et al. 2022; McGinley et al. 1980; Shafiuddin et al. 2026; Zhou et al. 2020). Beyond occupying this niche, *C. acnes* exerts metabolic activities that modify the local environment. *C. acnes* ferments host-derived substrates into short-chain fatty acids (SCFAs): acetate, propionate, and butyrate (Huang et al. 2022). These SCFAs acidify the local environment, enhance skin barrier (Bolla et al. 2020), inhibit competing microbes (Nakamura et al. 2020), and modulate the skin’s immunogenic response (Sanford et al. 2019). Furthermore, the bacterium secretes extracellular lipases that hydrolyze sebum triglycerides into free fatty acids, essential for its growth (Higaki 2003; Holland et al. 2010; Nakase et al. 2022; Whiteside and Voss 1973). These activities suggest that *C. acnes* not only adapts to but also actively shapes the follicular niche.

*C. acnes* abundance closely tracks sebaceous gland (SG) activity across the lifespan, a process primarily governed by androgens (Leyden 1995). *C. acnes* colonization is minimal during infancy (Capone et al. 2011; Dominguez-Bello et al. 2010), but increases during puberty, coinciding with androgen-stimulated sebum production (Capone et al. 2011; Dwyer and Scharschmidt 2022; Schneider et al. 2023). This relationship reverses with aging, where reduced sebum output, impaired sebocyte turnover, and eventual glandular atrophy limit lipid availability and reduce *C. acnes* colonization (Challa et al. 2025; Hou et al. 2022; Pochi et al. 1979). These observations highlight the tight coupling between SG function and *C. acnes* abundance but have largely been interpreted as a unidirectional host-driven process.

Emerging evidence suggests *C. acnes* can influence host lipid metabolism by stimulating lipid production in keratinocytes (Almoughrabie et al. 2023) and hamster sebocytes (Iinuma et al. 2009). However, its role in regulating sebogenesis and late-stage differentiation of human sebocytes remains unclear. This distinction is critical, as sebum secretion is a specialized development program in which sebocytes transition from an undifferentiated state to a terminal, lipid-filled phenotype, followed by nuclear size changes and degradation, accompanied by cellular enlargement, ultimately releasing their lipid content through holocrine secretion (Plewig and Christophers 1974; Tosti 1974; Wróbel et al. 2003; Zouboulis 2017; Zouboulis et al. 2016; Zouboulis et al. 1994). Hence, while microbial influence on lipogenesis has been suggested, *C. acnes’* role in driving human sebocyte differentiation and holocrine secretion remains unknown.

Given *C. acnes’* ecological dependence on sebum-derived lipids, we hypothesized *C. acnes* metabolites actively modulate sebocyte function – stimulating lipogenesis and promoting differentiation towards holocrine secretion. To test this, we examined how *C. acnes-*derived metabolites affect primary human sebocytes and the mechanisms through which they regulate lipid synthesis and maturation.

## RESULTS

### *C. acnes* secretome induces lipogenesis in primary human sebocytes

To test whether *C. acnes* actively modulates sebocyte lipid synthesis, we treated primary human sebocytes with cell-free supernatant (CFS) of *C. acnes* and quantified lipid droplet (LD) number per cell, LD area (individual LD size), LD intensity, nuclear count, and nuclear area via an automated image-based assay (Figure 1A).

**Figure 1.**
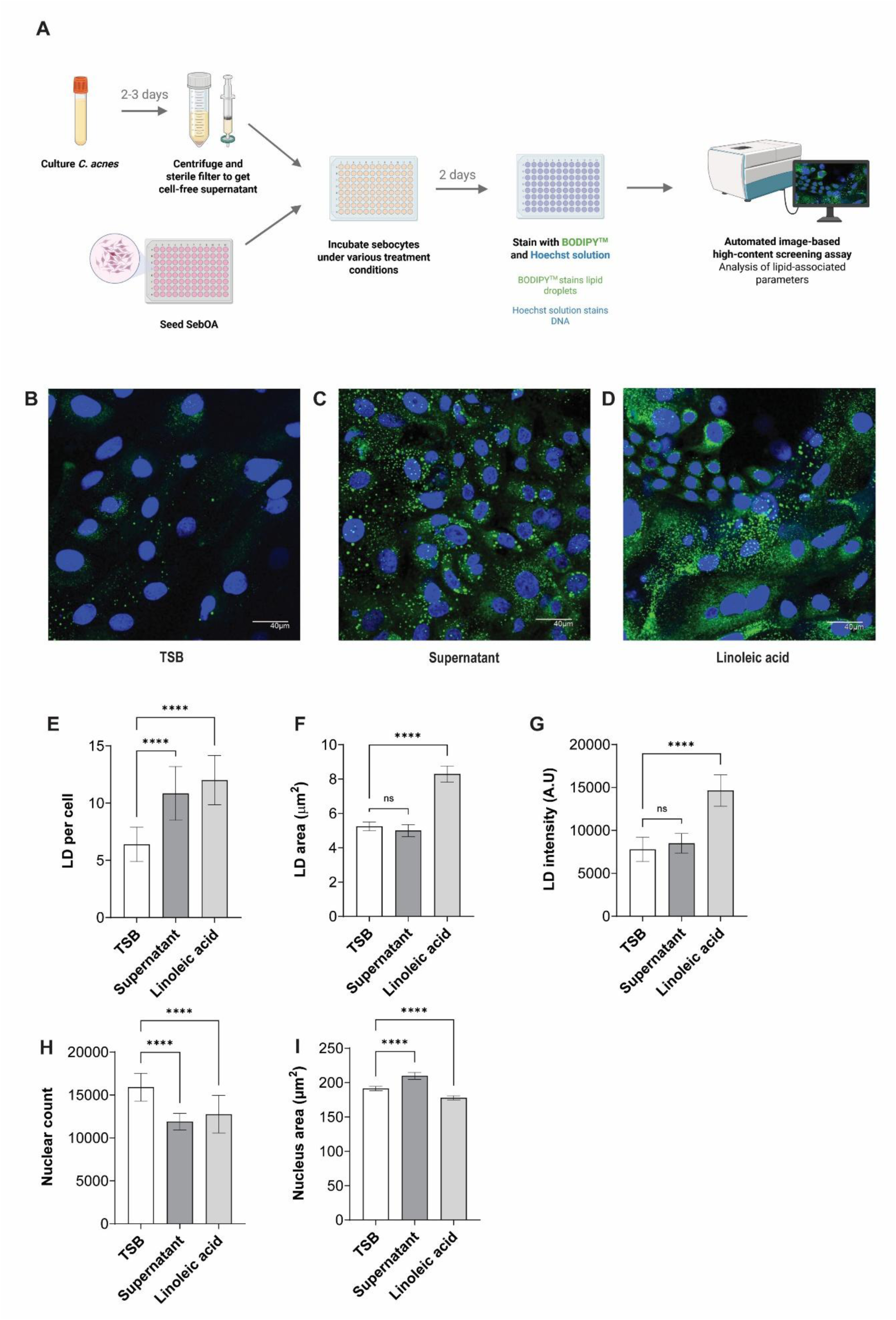
Cell-free supernatant of *C. acnes* leads to increased lipid production in sebocytes. **(A)** Experimental workflow schematic. Micrographs of sebocytes cultured for 48h with **(B)** 25% (v/v) fresh TSB, **(C)** 25% (v/v) Day 3 *C. acnes* CFS, or **(D)** 60µM linoleic acid. Hoechst (blue, nuclei) and BODIPY (green, LD). Scale bar, 40 µm. Quantification of **(E-G)** LD and **(H-I)** nuclear properties in sebocytes after 48h treatment with 25% (v/v) TSB (*n* = 16-17), 25% (v/v) Day 3 CFS (*n* = 24-25), or 60µM linoleic acid (*n* = 16-17). Error bars: mean ± SD. ANOVA: \**P*<0.05, \*\**P*<0.01, \*\*\**P*<0.001, and \*\*\*\**P*<0.0001. CFS, cell-free supernatant; LD, lipid droplet; SebOA, primary human sebocytes; TSB, tryptic soy broth.

CFS treatment increased lipogenesis, as evidenced by the significant increase in LD number per cell compared to the negative control (p < 0.0001; Figure 1B-E), without altering LD area and intensity (Figure 1F-G). This indicates that *C. acnes* secretome primarily increased LD abundance without altering lipid packing density within individual LDs. Notably, CFS treatment also reduced nuclear count and increased nuclear area (p < 0.0001; Figure 1H-I). This pattern is consistent with a reduction in intact cell numbers accompanied by morphological changes during late differentiation stages. Given that sebocytes undergo holocrine secretion involving terminal differentiation, followed by disintegration of lipid-rich cells to release lipids *in situ*, these observations indicate that *C. acnes* CFS not only stimulates lipogenesis but may also promote holocrine secretion of lipids from sebocytes. Together, these results demonstrate that *C. acnes* secretome drives lipogenesis in human sebocytes and is consistent with holocrine secretion induction.

### SCFAs from *C. acnes* drives lipogenic response

To identify the *C. acnes* metabolites responsible for lipogenesis, we analysed the *C. acnes* CFS using gas chromatography with flame ionisation detection (GC-FID), which identified propionate and acetate as the predominant metabolites in the secretome (Figure 2A-B). We focused on these SCFAs as they represent major metabolic end-products of *C. acnes* fermentation, with established roles in modulating host cell metabolism (Kao et al. 2021; Sanford et al. 2019).

**Figure 2.**
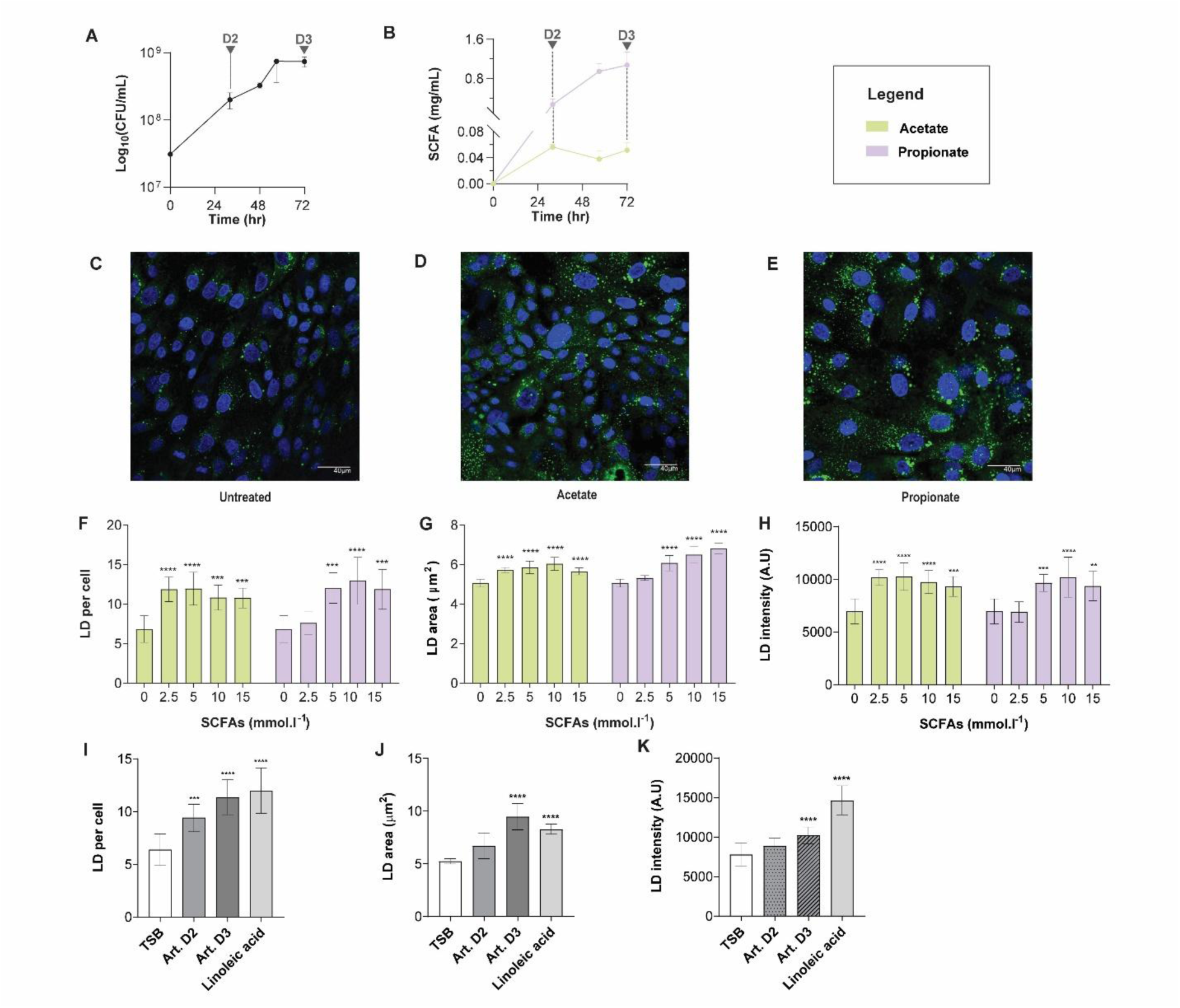
Acetate and propionate trigger lipid production in sebocytes. **(A)** *C. acnes* growth curve, and **(B)** acetate and propionate quantification in CFS over 72h (n = 3). Micrographs of sebocytes treated with **(C)** 25% (v/v) TSB, **(D)** 5mM acetate, or **(E)** 5mM propionate for 48h (*n* = 8). Hoechst (blue, nuclei) and BODIPY (green, LD). Scale bar, 40 µm. Quantification of LD properties in sebocytes treated for 48h with **(F-H)** increasing acetate and propionate concentrations, **(I-K)** 25% (v/v) TSB, 60µM linoleic acid, or 25% (v/v) artificial supernatants mimicking Day 2/3 *C. acnes* supernatants (*n* = 8-17). Error bars: mean ± SD. ANOVA: \**P*<0.05, \*\**P*<0.01, \*\*\**P*<0.001, and \*\*\*\**P*<0.0001. Art., artificial supernatant; CFS, cell-free supernatant; LD, lipid droplet; SCFA, short-chain fatty acid; TSB, tryptic soy broth.

We tested if these SCFAs are sufficient to recapitulate the lipogenic response of the *C. acnes* secretome. Sebocytes treated with 5mM acetate or propionate significantly increased LD number, area, and intensity (p < 0.0001, except propionate LD number and intensity p < 0.001; Figure 2 C-H), indicating enhanced lipid accumulation. Notably, propionate showed a dose-dependent increase in LD area (Figure 2F-H), suggesting a stronger effect on total lipid content than acetate. To validate that SCFAs account for the native secretome activity, we generated an artificial CFS by supplementing TSB with acetate and propionate at concentrations matching Day 2 (D2) and Day 3 (D3) *C. acnes* supernatants. This reconstituted mixture reproduced the lipogenic phenotype observed with native CFS (Art. D3 supernatant: p < 0.0001; Figure 2I-K), demonstrating that these SCFAs are sufficient to drive lipogenesis.

Collectively, these results identify SCFAs as the principal bioactive components of the *C. acnes* secretome associated with sebocyte lipogenesis and establish propionate as the principal metabolite that drives sebogenesis.

### Propionate reprograms central carbon metabolism in sebocytes

To investigate mechanisms underlying *C. acnes-*induced lipogenesis, we performed microarray-based pathway analysis in sebocytes treated with propionate (15mM), acetate (15mM), or *C. acnes* CFS (25% v/v D3). Gene expression changes were mapped across central carbon metabolism pathways to determine whether SCFAs reorganize sebocyte carbon flux. Propionate, but not acetate, induced coordinated reprogramming of central carbon metabolism, consistent with redistribution of carbon inputs towards lipogenic output. A similar overall pattern was observed in sebocytes treated with *C. acnes* CFS, further supporting propionate as the key metabolic driver of bacterial secretome.

At the level of carbon acquisition, propionate increased glucose transporter expression (Figure 3A), indicating enhanced carbon uptake. With glycolysis, a hexokinase isoform shift from HK2 to HK1 (Figure 3B) suggested a transition away from a proliferative glycolytic state towards intracellular carbon retention (Wilson 2003). Consistently, fructose-1,6-biphosphatase 1 (FBP1) upregulation with glucose-6-phosphatase (G6PC) unchanged (Figure 3B) indicates activation of a partial gluconeogenic programme that retains carbon at glucose-6-phosphate and increases substrate availability for the pentose phosphate pathway (PPP). Upregulation of the rate-limiting PPP enzyme, glucose-6-phosphate dehydrogenase (G6PD) (Figure 3C), enables NADPH production for reductive lipid biosynthesis (Schmidt et al. 2025).

**Figure 3.**
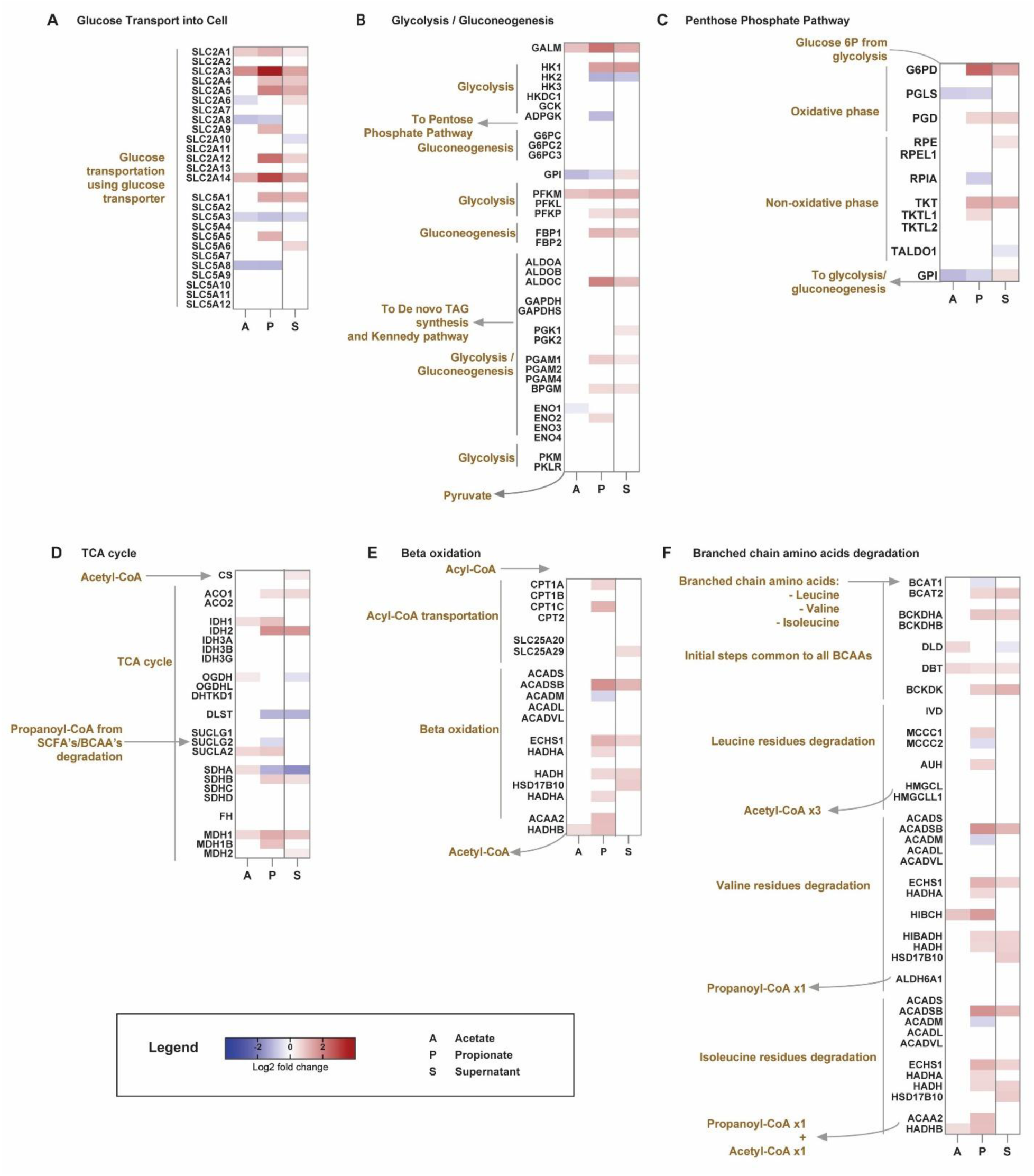
Propionate reprograms central carbon and mitochondrial metabolism in sebocytes to support energy production for lipogenesis. Heatmap showing log2 fold-change gene expression for **(A)** glucose transport, **(B)** glycolysis and gluconeogenesis, **(C)** pentose phosphate pathway, **(D)** TCA cycle, **(E)** mitochondrial beta oxidation, and **(F)** BCAA in sebocytes treated with 15mM acetate, 15mM propionate, or 25% (v/v) Day 3 *C. acnes* CFS (n = 3). Log2 fold-changes are relative to matched controls (untreated for acetate/propionate; TSB for CFS). Red, upregulation; blue, downregulation; white, P > 0.05. BCAA, branched-chain amino acid; CFS, cell-free supernatant; CoA, coenzyme A; Glucose 6P, glucose 6-phosphate; SCFA, short-chain fatty acid; TAG, triacylglycerol; TCA, tricarboxylic.

Propionate further reshaped mitochondrial metabolism through selective tricarboxylic acid (TCA) cycle remodelling, consistent with biosynthetic precursor expansion. The E2 subunit of the α-ketoglutarate dehydrogenase (DLST) expression was reduced upon propionate treatment, accompanied by an isoform shift in succinyl-CoA ligase (SUCLA2 up, SUCLG2 down), suggesting promotion of ATP production to meet the energy need (Miller et al. 2011), and discordant regulation of complex II (SDHB up, SDHA down), indicating active metabolic flux reprogramming (Figure 3D). In parallel, aconitase 1 (ACO1), isocitrate dehydrogenases (IDH1, IDH2), and malate dehydrogenases (MDH1, MDH1B) were upregulated, suggesting enhanced citrate/isocitrate and malate-linked pathways, supporting cytosolic acetyl-CoA and NADPH generation (Figure 3D). Despite propionate’s ability to enter the TCA cycle via succinyl-CoA, the canonical propionate catabolic pathway was not coordinately induced, with only a modest upregulation of propionyl-CoA carboxylase alpha subunit (PCCA, Log2FC = 1.1) and no significant change in PCCB or methylmalonyl-CoA mutase (MUT) (Figure 3D). In parallel, branched-chain amino acid catabolism was induced (Figure 3F), with leucine, valine, and isoleucine contributing to acetyl-CoA and propionyl-CoA pools. Furthermore, mitochondrial beta-oxidation was upregulated (Figure 3E), including strong induction of short-chain acyl-CoA transporters, suggesting enhanced shuttling of beta-oxidation-derived short-chain intermediates for further metabolic processing. Together, these changes indicate convergence of glucose, amino acid, and lipid-derived carbons into the shared acetyl-CoA and propionyl-CoA pools, while simultaneously maintaining ATP production to meet the energetic demands for lipogenesis.

This metabolic convergence is reinforced at the level of cytosolic acetyl-CoA generation. ATP-citrate lyase (ACLY) and acetyl-CoA synthetase (ACSS2) were upregulated under propionate treatment, indicating expansion of parallel pathways that generate cytosolic acetyl-CoA. ACSS2 enables conversion of free acetate generated from beta-oxidation and other processes to acetyl-CoA. Together, these data indicate that propionate-treated sebocytes integrate multiple carbon inputs into a shared acetyl-CoA pool while expanding NADPH and energy supply to support anabolic lipid metabolism.

### Propionate activates *de novo* lipid synthesis and lipid assembly pathways

Having established that propionate reorganises central carbon metabolism, we investigated how this reprogramming drives lipid synthesis. Propionate strongly induced core triacylglycerol (TAG) synthesis pathways through fatty acid synthase (FASN) and acetyl-CoA carboxylase (ACACA) upregulation, generating malonyl-CoA for fatty acid elongation (Figure 4A). Notably, propionate-induced FASN expression exceeded that of acetate or CFS, strongly suggesting that propionate actively drives the transcriptional fatty acid synthesis program in sebocytes.

**Figure 4.**
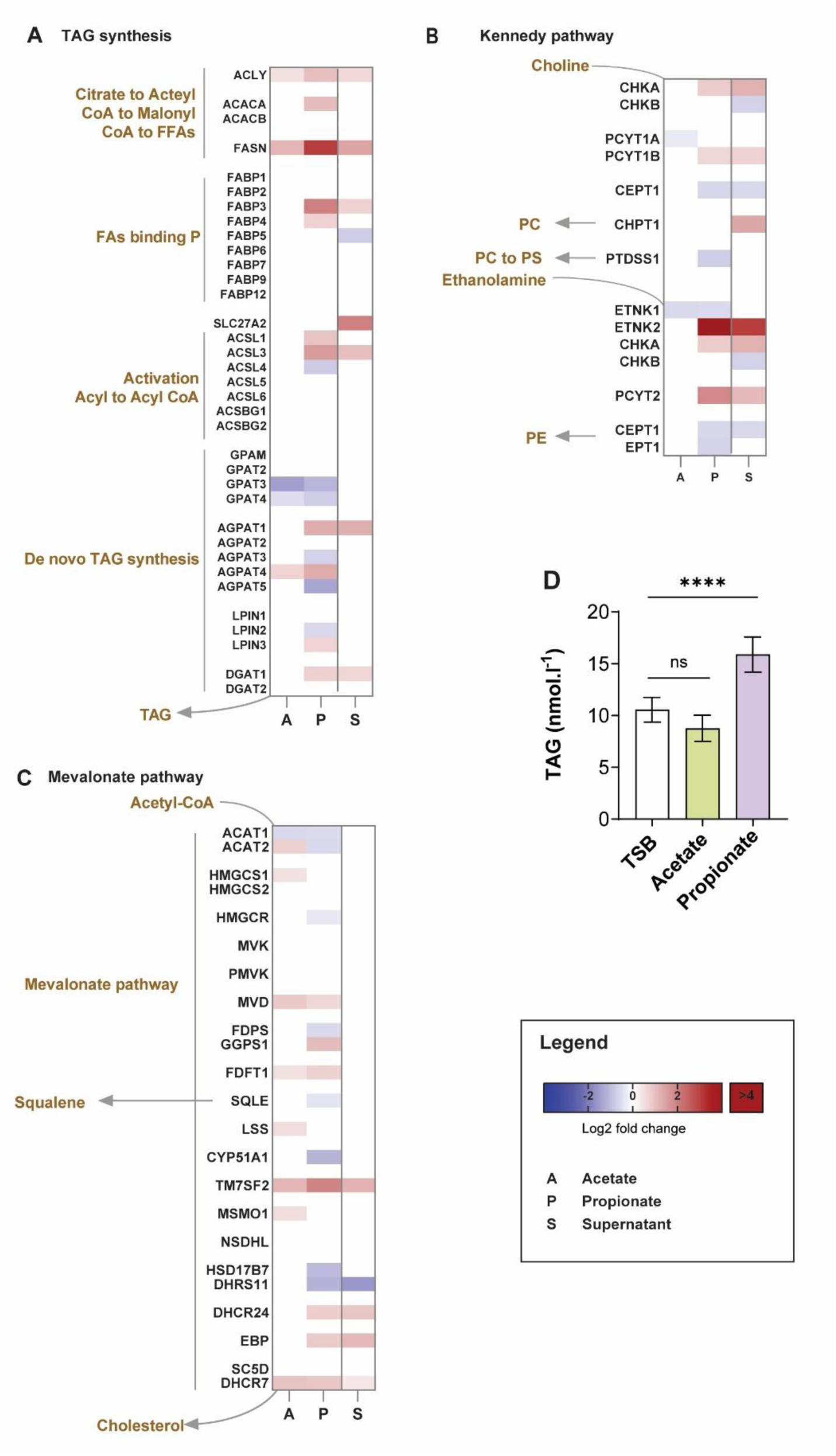
Propionate enhances lipid biosynthesis pathways in sebocytes. Heatmap showing log2 fold-change gene expression for **(A)** TAG synthesis, **(B)** Kennedy pathway, and **(C)** mevalonate pathway in sebocytes treated with 15mM acetate, 15mM propionate, or 25% (v/v) Day 3 *C. acnes* CFS (n = 3). Log2 fold-changes are relative to matched controls (untreated for acetate/propionate; TSB for CFS). Red, upregulation; blue, downregulation; white, P > 0.05. **(D)** TAG released by sebocytes after 48hr treatment with 25% (v/v) TSB, 15mM acetate, or 15mM propionate (n = 7-8). Error bars: mean ± SD. ANOVA: \**P*<0.05, \*\**P*<0.01, \*\*\**P*<0.001, and \*\*\*\**P*<0.0001. CFS, cell-free supernatant; CoA, coenzyme A; FAs, fatty acids; FFAs, free fatty acids; PE, phosphatidylethanolamine; PC, phosphatidylcholine; PS, phosphatidylserine; TAG, triacylglycerol; TSB, tryptic soy broth.

Downstream of fatty acid synthesis, propionate promoted lipid assembly remodelling pathways, channelling newly synthesized fatty acids into complex lipids. Isoform-specific changes in TAG synthesis enzymes were observed, including regulation of 1-acylglycerol-3-phosphate O-acyltransferases (AGPAT1/4 up, AGPAT3/5 down) (Figure 4A), pointing to a reprogramming of acyl-chain selectivity in glycerolipid biosynthesis (Karagiota et al. 2022). The terminal enzyme diacylglycerol O-acyltransferase (DGAT1) showed comparatively modest transcriptional changes, indicating final TAG assembly may be regulated primarily by substrate availability or post-transcriptional mechanisms.

Within the Kennedy pathway, strong ethanolamine kinase 2 (ETNK2) induction indicates enhanced phosphotidylethonolamine biosynthesis (Figure 4B), consistent with membrane remodelling required for LD formation. Furthermore, aldolase C (ALDOC) upregulation without corresponding glyceraldehyde-phosphate dehydrogenase (GAPDH) induction suggests triose phosphate pool expansion, increasing glycerol-3-phosphate availability required for TAG backbone synthesis (Figure 3B). These changes ensure fatty acyl chains and glycerol backbones are supplied for TAG lipid assembly.

The mevalonate-cholesterol pathway, on the other hand, exhibited a selective response. While mevalonate diphosphate decarboxylase (MVD) and transmembrane 7 superfamily member 2 (TM7SF2) were upregulated, the rate-limiting HMG-CoA reductase (HMGCR) showed minimal change (Figure 4C), suggesting increased flux through the distal pathway is driven by substrate availability from the acetyl-CoA pool rather than transcriptional activation. Consequently, squalene, a major sebum lipid component and an intermediate of this pathway, may accumulate as a result of enhanced mevalonate flux.

To validate the propionate-induced lipogenic program, we investigated whether SCFA treatment alters the secreted lipid composition. Propionate treatment significantly increased secreted TAG levels compared to the TSB control (p < 0.0001; Figure 4D), indicating enhanced lipid biosynthesis and secretion. In contrast, acetate had a comparatively modest effect, highlighting a functional divergence between SCFAs.

Together, these findings demonstrate that this metabolic reprogramming is tightly coupled with lipid synthesis and assembly pathways, culminating in a propionate-driven lipogenic state.

### Propionate induces a transcriptional programme consistent with sebocyte maturation and terminal differentiation

Beyond lipid synthesis, we investigated whether propionate promotes the sebocyte maturation required for holocrine secretion. To assess the sebocyte differentiation stages, we leveraged established markers from published high-resolution maps of sebocyte differentiation (Durgin et al. 2026; Düz et al. 2026; Schmidt et al. 2025; Schmidt et al. 2024; Yaba et al. 2024). Differential expression analysis revealed gene induction associated with multiple sebocyte differentiation stages, spanning progenitor (Seb-B), lipogenic (Seb-1/2), and terminal (Seb-3) states (Figure 5). This suggests that propionate drives sebocytes preferentially toward terminal differentiation rather than progenitor expansion.

**Figure 5.**
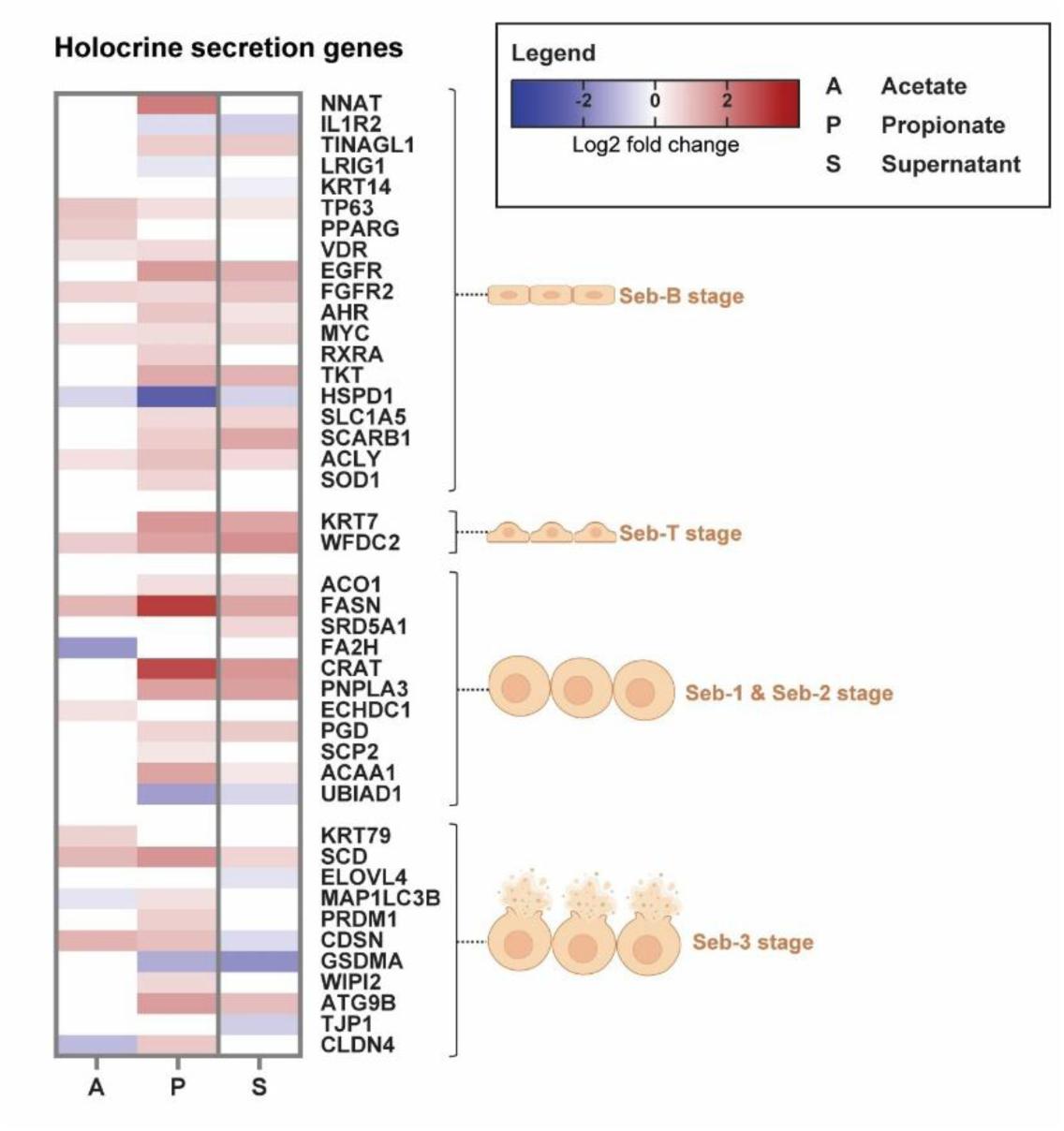
Propionate induces differentiation cascade in sebocytes leading to holocrine secretion. Heatmap showing log2 fold-change gene expression across the sebocyte maturation stages (Seb-B, Seb-T, Seb-1/2, Seb-3) in sebocytes treated with 15mM acetate, 15mM propionate, or 25% (v/v) Day 3 *C. acnes* CFS (n = 3). Log2 fold-changes are relative to matched controls (untreated for acetate/propionate; TSB for CFS). Red, upregulation; blue, downregulation; white, P > 0.05. CFS, cell-free supernatant; TSB, tryptic soy broth.

Propionate promotes progenitor sebocyte pool expansion, evidenced by the progenitor markers upregulation, neuronatin (NNAT) and epidermal growth factor receptor (EGFR). The induction of EGFR suggests that the cells are maintained in a pro-differentiation stage. During active lipid production stages (Seb-1/2), FASN and patatin-like phospholipase domain-containing 3 (PNPLA3) are upregulated, indicating that the cells are actively synthesizing and organizing lipid into storage droplets, consistent with the lipogenic program described in the preceding sections. As the cell enters Seb-3 to prepare for terminal maturation and lysis, autophagy-related protein autophagy-related 9B (ATG9B) and PR domain-containing protein 1(PRDM1) were upregulated, suggesting that propionate induces autophagy and terminal differentiation in sebocytes.

Furthermore, tight junction marker claudin 4 (CLDN4) was expressed to form a barrier in the uppermost nucleated sebocytes to prepare for holocrine secretion. Unlike propionate, acetate and *C. acnes* CFS did not show transcriptional activation across the full differentiation cascade.

Taken together, these transcriptomic signatures, combined with the lipogenic effects, demonstrate that propionate induces lipogenesis and sebocyte maturation towards holocrine secretion.

## DISCUSSION

The classical model of the pilosebaceous unit assumes a unidirectional relationship, where androgen-driven sebum production passively supports *C. acnes* colonization. Our findings interrogate this model (Figure 6), demonstrating that *C. acnes* SCFAs actively reprogramme sebocyte metabolism to drive lipogenesis, remodel sebum composition, and trigger holocrine secretion, effectively constructing the niche it occupies.

**Figure 6.**
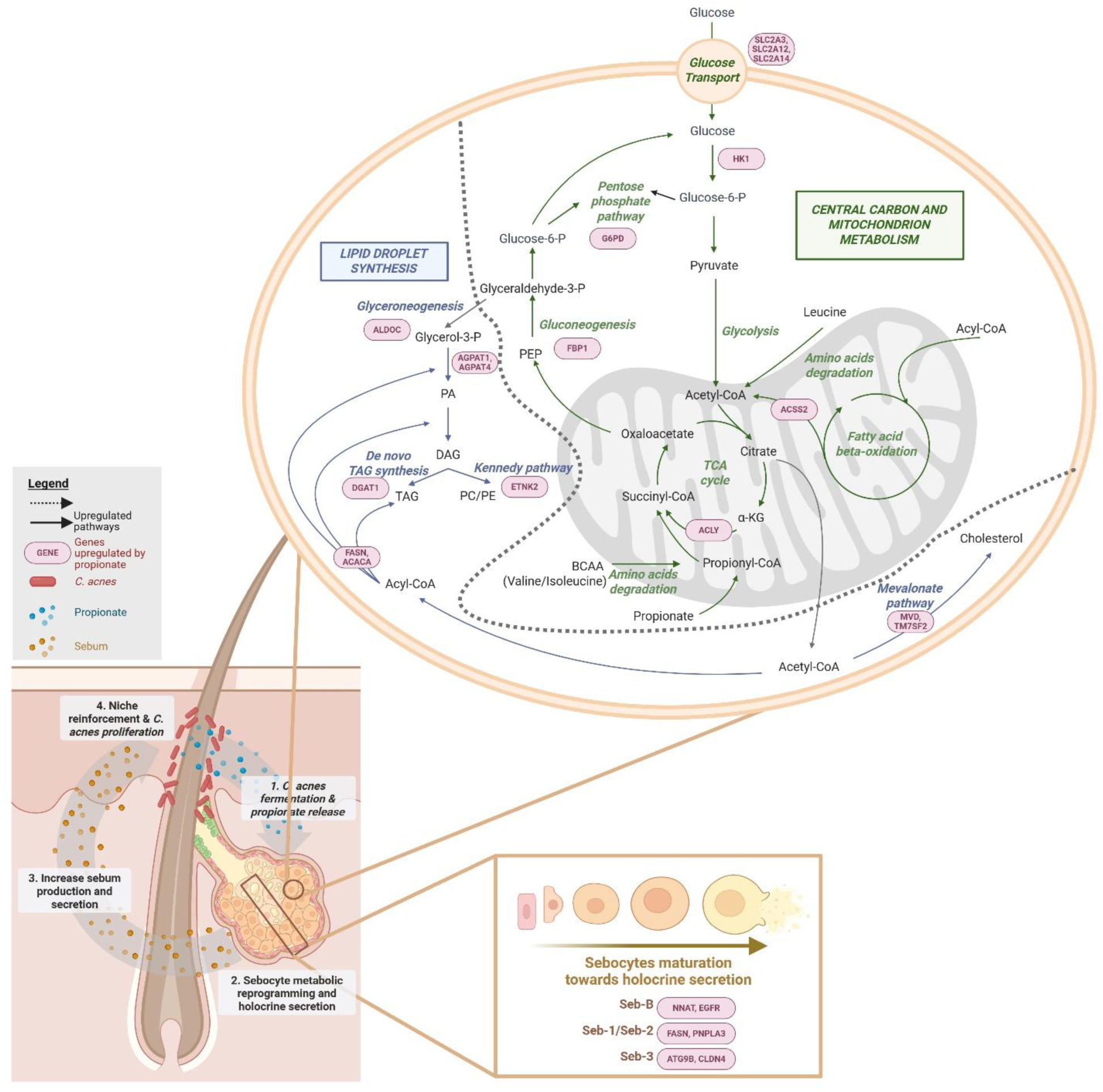
Proposed model of *C. acnes*-mediated metabolic and cellular reprogramming of sebocytes. *C. acnes-*derived propionate initiated a host-microbe metabolic feedback loop. At the cellular level, propionate upregulates key genes (in red boxes) in sebocytes involved in lipid droplet synthesis, central carbon and mitochondrial metabolism, and sebocyte maturation. These transcriptional changes collectively drive sebocytes metabolic reprogramming and accelerate holocrine secretion, resulting in increased sebum production and secretion into the follicular environment. The lipid-enriched follicular niche then reinforces *C. acnes* colonization and proliferation, perpetuating the feedback loop. PA, phosphatidic acid; DAG, diacylglycerol; TAG, triacylglycerol; PC, phosphatidylcholine; PE, phosphotidylethanolamine; PEP, phosphoenolpyruvate; Glucose 6-P, glucose 6-phosphate; Glyceraldehyde 3-P, glyceraldehyde 3-phosphate; TCA cycle, tricarboxylic acid cycle; α-KG, alpha-ketoglutarate; BCAA, branched-chain amino acids; CoA, coenzyme A.

We first show that *C. acnes* secretome induces lipid production in human sebocytes (Figure 1). While prior studies in hamster sebocytes showed that *C. acnes* can enhance lipogenesis (Iinuma et al. 2009), specific microbial mediators remained elusive. Through metabolic profiling, we identified acetate and propionate as the dominant SCFAs produced by *C. acnes*, with propionate as the primary lipogenic driver (Figure 2). While *C*. *acnes*-derived SCFAs have been shown to drive lipogenesis in keratinocytes via PPARα activation (Almoughrabie et al. 2023), our findings extend this biology to the SG – the skin’s primary lipid producer – where propionate remodels the central carbon metabolism towards the generation of energy and biosynthetic precursors for lipid assembly (Figure 3).

Importantly, these findings provide a mechanistic extension to a proposed model of sebaceous gland metabolism (Schmidt et al. 2025). While the proposed model has reconstructed the metabolic programme underlying sebocyte differentiation from steady-state transcriptomic trajectories, identifying coordinated glucose uptake, PPP activation and TCA-cycle remodelling as hallmarks of lipogenic sebocytes, it did not identify the upstream signal initiating this programme. Our data shows that microbial-derived propionate is a physiological trigger for this metabolic programme. Propionate recapitulated the induction of glucose transporters, G6PD-dependent redirection into the PPP and citrate-generating TCA remodelling described in this model, accounting for much of the transcriptional response observed with the *C. acnes* secretome. We note that the canonical propionate catabolic pathway that enable propionate to enter TCA through conversion of Succinyl-CoA is not significantly enriched in the transcriptome (Fig. 3D). Therefore, it is unclear if propionate acts as a signal or used as a substrate for energy metabolism. However, these findings identify propionate as a microbial metabolite capable of initiating a host metabolic programme leading to sebogenesis.

A second key finding is that propionate remodels sebum lipid composition, specifically increasing TAG accumulation (Figure 4). This shift was accompanied by isoform-specific reprogramming of AGPAT expression, consistent with the redirection of acyl-chain selectivity away from organelle membrane phospholipid synthesis and towards glycerolipid storage (Karagiota et al. 2022). Sebum TAGs are the primary substrates hydrolysed by *C. acnes* lipases to generate free fatty acids that sustain bacterial growth (Li et al. 2025). This establishes a feed-forward loop, where propionate drives TAG synthesis, fuelling *C. acnes* proliferation and further propionate production. This feed-forward architecture is the biochemical basis of niche construction.

A third, and most striking finding is that propionate induces sebocytes differentiation cascade towards holocrine secretion (Figure 5) – a process traditionally considered exclusively driven by endogenous host-hormonal and lipid signals (Wróbel et al. 2003; Zhang et al. 2018). Our data show that propionate activates a transcriptional program, encompassing autophagy-related genes (ATG9B, MAP1LC3B, WIPI2) and late-stage Seb-3 genes, consistent with holocrine priming. While our single-timepoint transcriptomic analysis in a 2D-system cannot resolve the temporal differentiation progression, concurrent enrichment of stage-specific markers identifies a clear microbial trigger for sebocyte differentiation (Figure 6).

Propionate, however, is not solely a sebocyte effector. Across the skin tissue, it engages multiple cell types through context-dependent mechanisms converging on coherent physiological function. In keratinocytes, propionate induces autophagy via MAPK activation and mitochondrial damage (Megyeri et al. 2018), and modulates inflammatory responses through HDAC activity relevant to acne pathogenesis (Sanford et al. 2016). It also alleviates atopic dermatitis itch via TRP channel modulation and mTORC2 inhibition (Kim et al. 2024; Xu et al. 2024). Collectively, these observations position propionate as a pleiotropic signalling metabolite that coordinates multiple dimensions of skin physiology, including lipid production, barrier integrity, pigmentation, immune tone, and sensory signalling. This convergence reflects that *C. acnes* and its metabolite, propionate, are best understood as regulators of not only the skin biochemical environment but also tissue function.

The implications of propionate-driven skin niche modulation extend beyond *C. acnes*. Sebum is a key ecological filter shaping the skin microbiome (Grice and Segre 2011), but it has generally been treated as a static host-derived substrate. Our findings challenge this view, positioning sebum as a dynamically regulated output of host-microbe interactions. The propionate-driven increase in TAG is particularly consequential for lipophilic organisms like *Malassezia*, which is strictly lipid-dependent and expresses lipases that hydrolyse TAGs to release free fatty acids (Ro and Dawson 2005; Sommer et al. 2015). This increase in TAGs could support fungal overgrowth in contexts like truncal acne, where *Malassezia* could exacerbate follicular inflammation via free fatty acid release (Dawson 2007). Concurrently, *C. acnes*-derived propionate has been shown to suppress *Malassezia resticta* and *Staphylococcus epidermidis* in dandruff (Rasheedkhan Regina et al. 2024)^1^, and *Staphylococcus aureus* (Jeong et al. 2019), suggesting a regulatory mechanism influencing the skin microbiome. Nevertheless, the extent to which *C. acnes*-driven sebum remodelling alters the competitive landscape for the broader community remains an open question, but our data suggest that the follicular niche cannot be understood without accounting for *C. acnes* as an active regulator of its biochemical environment.

In conclusion, our findings position *C. acnes* as an active modifier of the sebaceous niche, with propionate identified as a key microbial metabolite reprogramming host central metabolism and differentiation. Sebocyte function is not solely governed by endogenous hormonal cues but is actively modulated by microbiome-derived signals. Collectively, this study redefines the *C. acnes*-SG relationship, establishing a mechanistic link between microbial metabolism and host tissue function.

## MATERIALS AND METHODS

### Bacteria cultures

*C. acnes* (ATCC-6919) was pre-cultured on tryptic soy agar (TSA; BioMérieux) for 5-7 days at 33°C. Single colonies were inoculated into snap-cap tubes (Greiner Bio-One) containing tryptic soy broth (TSB; BioMérieux) and incubated anaerobically for 5 days at 33°C. Each tube served as a biologically independent replicate. Cultures were standardized to OD_600_ = 0.05 ± 0.005 (Infinite-200 PRO, Tecan) before anaerobic incubation at 33°C for growth curves and CFS collection.

For growth curves, bacterial growth was monitored by OD_600_ and colony-forming units (CFUs) over 4 days. CFUs were determined from serial dilutions on TSA and incubated anaerobically at 33°C for 5 days.

For CFS, cultures were centrifuged at designated time points (4500rcf, 5 min), and supernatants were filtered (0.2µm).

### Sebocyte culture

G7F(-) media comprised DMEM glutamax and F12 glutamax (3:1 ratio; Gibco^TM^), supplemented with 5µg/mL insulin, 2nM tri-iodothyronine, 10ng/mL epidermal growth factor, 5µg/mL transferrin, 1µM isoproterenol, 180µM adenine hydrochloride (all Sigma-Aldrich), and 10% serum fetal clone II (Hyclone^TM^). G7F(+) media was identical, with 0.4µg/mL hydrocortisone (Sigma-Aldrich) and 1X antibiotic-antimycotic (Gibco^TM^) added.

Mitomycin-treated 3T3 fibroblasts (3T3m; Rheinwald and Green 1975; Todaro and Green 1963) were seeded in 175-cm² flasks (Thermo Scientific) in G7F(+) media as feeder cells (patent: FR3061205-A1). After 2h, SebOA primary sebocytes (Biopredic International, Batch: PEA033195-OC01) were co-cultured on the 3T3m layer until 70-80% confluency. For subsequent experiments, cells were harvested using trypsin (Gibco^TM^) and seeded in 96-well collagen-coated plates (Greiner Bio-one; 10,000 cells/well) or 6-well plates (Falcon^®^, 100,000 cells/well) in G7F(-) media until 80-90% confluency. Incubations were at 37°C with 5% CO_2_.

### Volatile fatty acid analysis

Supernatants were diluted 10-fold with 0.1% formic acid and analysed by GC-FID (GC-2010, Shimadzu) using a DB-FFAP column (30m x 0.25mm x 0.25µm; Agilent Technologies).

### Sebogenesis assays

Sebocytes were treated for 48h with: 25% (v/v) *C. acnes* supernatant; 25% (v/v) artificial supernatant mimicking D2 (0.13 mg/mL sodium acetate, 0.41 mg/mL sodium propionate) or D3 (0.74mg/mL sodium acetate, 1.3mg/mL sodium propionate) SCFA concentrations; 25%(v/v) TSB; or varying concentrations of linoleic acid, sodium acetate, or sodium propionate (all Sigma-Aldrich).

After incubation, cells were fixed in 10% neutral-buffered formalin (Sigma-Aldrich). LDs were stained with 5µg/mL BODIPY solution (Invitrogen) in Dulbecco’s Phosphate-Buffered Saline (DPBS; Gibco) containing 1% bovine serum albumin (Sigma-Aldrich). Nuclei were counterstained with Hoechst 33342 (1:2500 ratio; Thermo Scientific). High-content screening was performed using an ImageXpress Micro 4 system (16 images/well, 10x objective; Molecular Devices, version 6.2.3.733) and analyzed with the MetaXpress Transfluor module.

### Microarray analysis

Sebocytes were treated for 24h with 15mM sodium acetate, 15mM sodium propionate, or 25% (v/v) D3 *C. acnes* CFS. RNA was extracted using the RNeasy Plus Micro Kit (Qiagen) and quantified by NanoDrop (Thermo Scientific). Microarray was performed by Research Instruments Pte Ltd (Singapore). 100ng RNA was prepared using the GeneChip® WT PLUS Reagent kit and hybridized to Clariom^TM^ S Human Arrays (all Affymetrix). Arrays were washed and scanned on an Affymetrix Scanner 3000 7G using AGCC software.

### Triacylglycerol quantification

Sebocytes were treated for 48h with 25% (v/v) TSB, 15mM sodium acetate, or 15mM sodium propionate. TAG released by sebocytes was quantified using the TAG Assay Kit (Abcam) with colorimetric detection (λ_abs_=570nm; Infinite-200 PRO, Tecan).

### Statistical analysis

Analyses were performed using Prism v10.3.1 (GraphPad Software Inc.). Between-group comparisons were performed using the Mann-Whitney test, Kruskal-Wallis with Dunn’s post-hoc, or one-way ANOVA with Tukey’s correction. Microarray raw data (.CEL files) were SST-RMA normalized in TAC 4.0 (Affymetrix), and CD Genomics performed downstream biostatistical analyses to obtain differentially expressed genes.

## ETHICS STATEMENT

This study required no institutional approval or written consent, as it used cell lines that is commercially obtained

## DATA AVAILABILITY STATEMENT

The transcriptomic data generated in this study will be made available through a publicly accessible repository prior to publication.

## ACKNOWLEDGEMENTS

We thank Chloe Trotzier for providing primary sebocytes. We thank Tze Khee Chan for her technical expertise. The authors would like to acknowledge the support of the National Research Foundation and Ministry of Education Singapore under its Research Centre of Excellence Program to the Singapore Centre for Environmental Life Sciences Engineering (EDUN C33-62-036-V4). We thank Emma Pilling, PhD, and Marielle Romet, PhD (Santé Active Edition – Synergy Pharm) for medical writing assistance.

## AUTHOR CONTRIBUTION STATEMENT

Conceptualization: OSK, ASLT, VRR, SR, TC; Methodology: FB, TC, ASLT, EK; Formal Analysis: OSK, ADL, EK; Investigation: FB, TC, ASLT; Data Curation: ADL; Visualization: ADL; Writing-Review and Editing: OSK, ADL, ASLT, TC, VRR; Supervision: OSK, ASLT, TC; Funding acquisition: SR, TC

## DECLARATION OF GENERATIVE AI USE

During the preparation of this work, the author(s) used Gemini 3 to improve the grammar and syntax of the manuscript text. After using this tool/service, the author(s) reviewed and edited the content as needed and take(s) full responsibility for the content of the publication.

## Footnotes

1 Preprint data: Rasheedkhan Regina V, Chopra T, Weihao K, Cheruvalli S, Sabrina A, Fatimah Binte Jamal Mohamed H, et al. Decoding scalp health and microbiome dysbiosis in dandruff. 2024

